# Case-control analysis identifies shared properties of rare germline variation in cancer predisposing genes

**DOI:** 10.1101/162479

**Authors:** Mykyta Artomov, Vijai Joseph, Grace Tiao, Tinu Thomas, Kasmintan Schrader, Robert J. Klein, Adam Kiezun, Namrata Gupta, Lauren Margolin, Alexander J. Stratigos, Ivana Kim, Kristen Shannon, Leif W. Ellisen, Daniel Haber, Gad Getz, Hensin Tsao, Steven M. Lipkin, David Altshuler, Kenneth Offit, Mark J. Daly

## Abstract

Traditionally, genetic studies in cancer are focused on somatic mutations found in tumors and absent from the normal tissue. Identification of shared attributes in germline variation could aid discrimination of high-risk from likely benign mutations and narrow the search space for new cancer predisposing genes. Extraordinary progress made in analysis of common variation with GWAS methodology does not provide sufficient resolution to understand rare variation. To fulfil missing classification for rare germline variation we assembled datasets of whole exome sequences from >2,000 patients with different types of cancers: breast cancer, colon cancer and cutaneous and ocular melanomas matched to more than 7,000 non-cancer controls and analyzed germline variation in known cancer predisposing genes to identify common properties of disease associated mutations and new candidate cancer susceptibility genes. Lists of all cancer predisposing genes were divided into subclasses according to the mode of inheritance of the related cancer syndrome or contribution to known major cancer pathways. Out of all subclasses only genes linked to dominant syndromes presented significant rare germline variants enrichment in cases. Separate analysis of protein-truncating and missense variation in this subclass of genes confirmed significant prevalence of protein-truncating variants in cases only in loss-of-function tolerant genes (pLI<0.1), while ultra-rare missense mutations were significantly overrepresented in cases only in constrained genes (pLI>0.9). Taken together, our findings provide insights into the distribution and types of mutations underlying inherited cancer predisposition.

---

Discovery of over 100 germline predisposition genes in cancer have not only revolutionized identification of individuals and families at higher risk, but also provided novel mechanistic insights into the role of pathways in cancer development and helped in mitigating the risk using appropriate clinical management^1^. Common in cancer genetics approach involves studying kindred with multiple samples and searching for DNA variation segregated between affected and non-affected members of the family. However, segregating mutations could be uniquely observed in a given kindred and do not provide compelling information about their capacity to explain cancer cases outside of kindred of interest. Multiple cohort-based studies of inherited variation in cancer with GWAS methods reached great success in identifying low to moderate risk common mutations^2^. Understanding rare coding variation on population scale requires massive genome/exome sequencing data both for cases and controls. Only recently sufficient statistical power was gained to discover new cancer susceptibility genes using case-control analysis of rare germline variation^3^. However, systematic description of rare inherited variation architecture in cancer cases in comparison to control subjects has not been reported yet.

In order to identify shared properties of rare germline variation in cancer 866 patients diagnosed with early onset and/or familial history either of breast cancer (MIM: [114480]), colon cancer (MIM: [114500]), cutaneous melanoma (MIM: [155600]), ocular melanoma (MIM: [155720]), Li-Fraumeni syndrome (MIM: [151623]) with family history and/or early onset of disease (Supplementary materials, Supplementary Table 1) were recruited at MGH (Boston, USA), MEEI (Boston, USA), MSKCC (New York, USA), Andreas Sygros Hospital (Athens, Greece). Written informed consent was obtained from all individuals. Patients in this cohort were subjected to initial genetic screening (Supplementary Methods) and further identified as “selected cases”. We additionally included 1754 cancer samples with matching phenotypes from The Cancer Genome Atlas with no ascertainment for family history and age of onset. This cohort was identified as “unselected cases”. Control set of 24,612 samples was assembled from dbGAP whole exome studies of non-cancer phenotypes (Supplementary Table 1). Whole exome DNA sequences from all samples were aligned on a reference genome and processed through GATK Haplotype Caller^4–6^ variant discovery pipeline as a single batch. Genotypes in assembled dataset were then subjected to quality check on per variant and per individual level (Supplementary Materials). To ensure close ancestral matching, we performed principal component analysis (PCA; Supplementary Figure 1A). For reduction of heterogeneity due to diverse population admixture, only the largest cluster of samples representing predominantly European ancestry was further analyzed. Within European-ancestry samples we performed relatedness analysis and removed all duplicates and first-degree relatives (PI_HAT > 0.2), resulting in total 846 selected cases, 1496 unselected cases and 7924 matched controls included in the final dataset used for analysis. Further, examination of common synonymous variants (MAF>5%) revealed a null-distribution of the Fisher’s exact test (two-sided) statistic between cases and controls with genomic inflation factor λ=1.012 (Supplementary Figure 1B).

Results from *Zhang et al.*^7^ provided good reference to known cancer susceptibility genes (germline and somatic) clustering based on dominant/recessive nature of linked cancer syndromes and contribution to common cancer pathways (Supplementary Table 2). We examined cumulative burden of rare (minor allele count of less or equal to 10) variants in cases compared to controls within each set of genes. Only genes linked to dominant cancer disorders exhibited significant burden in both selected and unselected cases compared to controls. Isolated analysis of damaging missense and protein-truncating variants (PTVs) established the main role of the latter in observed association signal (Supplementary Table 3A-D). We also observed enrichment in DNA repair pathway list only in selected cases, to further investigate this signal we noted, that *BRCA1* (MIM: [113705]) and *TP53* (MIM: [191170]) are present in both this and autosomal dominant disorders gene lists. We removed these two genes from both lists and repeated analysis. Autosomal dominant genes remain highly significant and DNA repair pathway genes association signal is lost (Sup. Table 3 E-F). Interestingly, significant abundance of risk alleles was observed both in selected (Two-sided Fisher’s test p=5.6×10^−8^; OR=3.53; OR CI=2.26-5.39) and unselected cases (Two-sided Fisher’s test p=1.28×10^−5^; OR=2.49; OR CI=1.66-3.69), however, burden of risk alleles in selected group was greater than in unselected group (Figure 1A). Genes linked to breast cancer disorders carry substantial number of PTVs in controls, while genes linked to more severe phenotypes, like Li-Fraumeni syndrome (*TP53*), uveal and cutaneous melanomas show no or very low count of PTV carriers in control cohort (Supplementary Figure 2). Downstream examination of allele frequency spectrum for mutations driving this association signal affirmed significance of singleton burden (Two-sided Fisher’s test p=1.49×10^−8^, p=2.36×10^−5^; OR=5.25, OR=3.23; OR CI=2.99-9.01, OR CI=1.88-5.46; selected and unselected cases, respectively) while variants with minor allele count 2-10 did not show significant enrichment (Two-sided Fisher’s test p=0.1, p=0.07 selected and unselected cases, respectively). Considering overrepresentation of PTVs in cases it was feasible to test genes linked to dominant cancer disorders for loss-of-function intolerance. We used probability of loss-of-function intolerance (pLI) from ExAC database^8^ to separate genes into loss-of-function tolerant (pLI<0.1) and intolerant (pLI>0.9) groups (Supplementary Figure 3). Given that our case cohort does not have pediatric cancer patients we expectedly observed significant burden of singleton PTVs only in tolerant genes (Two-sided Fisher’s test p=1.5×10^−8^, OR=3.66, OR CI=2.03-6.36; p=3.0×10^−4^, OR=2.74, OR CI=1.57-4.65, selected and unselected cases respectively, Figure 1B-C), as expected for adult onset disorder. While constrained genes are depleted in protein-truncating variants in cancer, we sought to test whether missense mutations are uniformly distributed between constrained and tolerant genes linked to dominant cancer syndromes. We did not observe any enrichment in damaging missense variants among cases using minor allele count of 1 (MAF~1×10^−4^) and 1-10 (MAF~1×10^−4^ - 1×10^−3^) as a frequency cutoff (Figure 1 D-F). To examine this further, we used non-TCGA subset of ExAC^8^ database to keep for analysis only variants with MAF<2.3×10^−5^ (not present in ExAC and singletons in ExAC non-Finnish Europeans). Ultra-rare missense variant analysis revealed significant burden in selected cases driven by loss-of-function intolerant gene contribution (Two-sided Fisher’s test p=0.045, p=0.025; OR=1.26, OR=1.44; OR CI=1.00-1.58, OR CI=1.03-1.97; all and constrained autosomal dominant disorder genes, respectively; Figure 1 G-I). Previous analysis of cutaneous melanoma cohort used in this study identified *EBF3* (MIM: [607407]) as a new germline predisposition gene demonstrating tumor suppressor functional activity^3^. Interestingly, this gene has pLI=1 and carried ultra-rare missense variation in conserved protein domains, consistently with our observations above.

**Figure 1.**
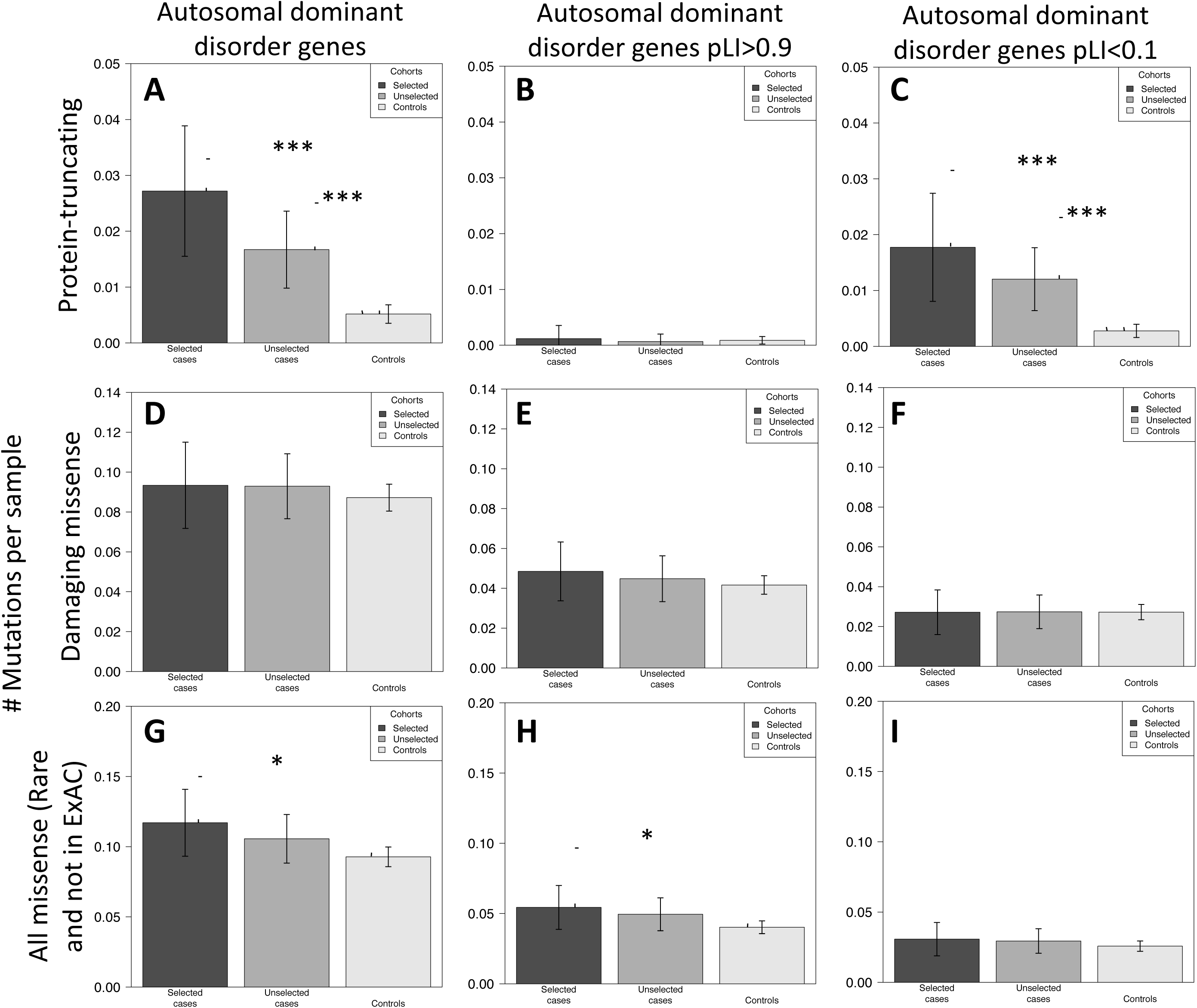
Mutational landscape overview. Mean mutation count per sample for protein truncating variants (MAC=1; MAF~1×10^−4^) (**A-C**); Damaging missense mutations (MAC=1; MAF~1×10^−4^) (**D-F**); Missense mutations that are not or rare (MAC=1; MAF=<2.3×10^−5^) in non-TCGA ExAC (**G-I**); estimated across all genes linked to autosomal dominant disorders (**A, D, G**); autosomal dominant disorders linked genes with pLI>0.9 (**B, E, H**); autosomal dominant disorders linked genes with pLI<0.1 (**C, F, I**); * - p<0.05; *** - p<0.001.

It is worth noting, however, that selected cases dataset was assembled by initial genetic screening of probands that satisfy NCCN genetic testing criteria^9^. If tested positive, they were not subsequently included in this study. Thus, genetically enriched cases have had more genetic screening and some diagnosed cases were removed before being entered in this study sample – likely attenuating the strength of association to the group with known autosomal dominant cancer predisposition genes.

Further, we subjected selected breast cancer cohort to independent analysis, as the largest cohort in our dataset. We kept only female samples for analysis, resulting in 354 breast cancer cases and 2171 controls subjected to downstream analysis. First, we analyzed burden of singleton PTVs in loss-of-function tolerant genes. *BRCA1* (p=1.34−10^−8^, 8/355 cases and 5/7924 controls) and *BRCA2* (p=6×10^−4^, 5/355 cases and 12/7924 controls) were two top genes in this analysis. Significance threshold was estimated using Bonferroni correction for number of genes tested – 0.05/7861 (3.34×10^−5^). Analysis of missense singletons in constraint genes did not return any significant genes likely due to power limitations.

Overall, observed germline variation in both selected and unselected cases in established cancer susceptibility genes is linked to dominant cancer disorders, majorly represented by PTVs and has ultra-low frequency in population (singletons). While we observed ultra-rare missense mutations enrichment in cases, proportion of cases explained by this type of variation is likely very small, though potentially providing additional genes contributing to cancer pathways. Understanding power limitations of our study and potential effects of imbalance between cancer cohort sizes, yet our results provide a reference point for allele frequencies and variation type for future search of new genes contributing to inherited cancer susceptibility through rare DNA variation. We expect that overall majority of cancer cases would be explained by sporadic somatic mutations and inherited polygenic risk (mostly driven by common DNA variation), however, analysis of enriched kindred is bound to ultra-rare variation which our results could significantly aid.

## WEB RESOURCES

OMIM, http://www.omim.org

ExAC database, http://exac.broadinstitute.org

Non-TCGA ExAC, ftp://ftp.broadinstitute.org/pub/ExAC_release/releaseQ.3.1/subsets

All case and control genotypes are publically available through dbGAP database.

## Notes

Authors declare no conflict of interests

